# Germline gene de-silencing by a transposon insertion is triggered by an altered landscape of local piRNA biogenesis

**DOI:** 10.1101/2020.06.26.173187

**Authors:** Danny E. Miller, Kelley Van Vaerenberghe, Angela Li, Emily K. Grantham, Celeste Cummings, Marilyn Barragan, Rhonda Egidy, Allison R. Scott, Kate Hall, Anoja Perera, William D. Gilliland, Justin P. Blumenstiel

**Affiliations:** Department of Pediatrics, Division of Medical Genetics, University of Washington, Seattle, Washington and Seattle Children’s Hospital, Seattle, WA 98105; Department of Ecology and Evolutionary Biology, University of Kansas, Lawrence, KS 66049; Stowers Institute for Medical Research, Kansas City, MO 64110; Department of Biological Sciences, DePaul University, Chicago, IL 60614

## Abstract

Transposable elements (TE) are selfish genetic elements that can cause harmful mutations. In *Drosophila*, it has been estimated that half of all spontaneous visible marker phenotypes are mutations caused by TE insertions. Because of the harm posed by TEs, eukaryotes have evolved systems of small RNA-based genome defense to limit transposition. However, as in all immune systems, there is a cost of autoimmunity and small RNA-based systems that silence TEs can inadvertently silence genes flanking TE insertions. In a screen for essential meiotic genes in *Drosophila melanogaster*, a truncated *Doc* retrotransposon within a neighboring gene was found to trigger the germline silencing of *ald*, the *Drosophila Mps1* homolog, a gene essential for meiosis. A subsequent screen for modifiers of this silencing identified a new insertion of a *Hobo* DNA transposon in the same neighboring gene. Here we describe how the original *Doc* insertion triggers flanking piRNA biogenesis and local gene silencing and how the additional *Hobo* insertion leads to de-silencing by reducing flanking piRNA biogenesis triggered by the original *Doc* insertion. These results support a model of TE-mediated silencing by piRNA biogenesis in *cis* that depends on local determinants of transcription. This may explain complex patterns of off-target gene silencing triggered by TEs within populations and in the laboratory. It also provides a mechanism of sign epistasis among TE insertions.

**Author Summary:** Transposable elements (TEs) are selfish DNA elements that can move through genomes and cause mutation. In some species, the vast majority of DNA is composed of this form of selfish DNA. Because TEs can be harmful, systems of genome immunity based on small RNA have evolved to limit the movement of TEs. However, like all systems of immunity, it can be challenging for the host to distinguish self from non-self. Thus, TE insertions occasionally cause the small RNA silencing machinery to turn off the expression of critical genes. The rules by which this inadvertent form of autoimmunity causes gene silencing are not well understood. In this article, we describe a phenomenon whereby a TE insertion, rather than silencing a nearby gene, rescues the silencing of a gene caused by another TE insertion. This reveals a mode of TE interaction *via* small RNA silencing that may be important for understanding how TEs exert their effects on gene expression in populations and across species.

## Introduction

Transposable elements (TE) are harmful selfish elements that can cause DNA damage, mutation, chromosome rearrangements, and sterility. Due to their capacity to cause mutation, it has been estimated that about half of the spontaneous mutations that cause visible phenotypes in *Drosophila* are caused by TE insertions [1]. Nonetheless, despite their harm, TEs can greatly proliferate in the genomes of sexually reproducing species [2]. A consequence of TE proliferation is that diverse systems of genome defense have evolved that limit transposition through DNA methylation, repressive chromatin, direct transcriptional repression, and small-RNA silencing. There is substantial cross-talk between these modes of genome defense. For example, small RNAs generated from harmful TE transcripts can silence TEs through cytoplasmic post-transcriptional silencing but also enter the nucleus to trigger DNA methylation and transcriptional repression [3,4].

In animals, small RNAs designated piwi-interacting RNAs (piRNAs) play a critical role in genome defense within reproductive tissues. piRNAs are derived from TE sequences recognized by the piRNA machinery and diverted from canonical mRNA processing into a piRNA generating pathway. By shunting TE transcripts toward piRNA biogenesis, the host is able to generate a pool of antisense piRNAs that repress TEs throughout the genome. Interestingly, like other systems of immunity, genomic immunity can be costly when the distinction between self and non-self is disrupted. For example, in *Arabidopsis thaliana*, selection can act against DNA-methylated TE insertions that reduce the expression of flanking genes [5]. Off-target gene silencing by systems of genome defense, and subsequent selective effects, has been observed in a variety of organisms [6–17]. However, genic silencing by flanking TEs is hardly universal within a genome. For example, in maize, the capacity to trigger the formation of flanking heterochromatin can vary significantly among TE families [13]. The cause of this variation is poorly understood.

Studies in *Drosophila*, where DNA methylation is absent and piRNAs are the primary line of defense against TEs, show that TE insertions can trigger the spreading of heterochromatin and transcriptional silencing of genes [16–21]. In the germline, TE insertions also have the capacity to trigger the production of piRNAs from flanking sequences [22–25]. The mechanism of flanking piRNA biogenesis that spreads from a TE insertion can be explained by a general model whereby Piwi-piRNA complexes target nascent TE transcripts [26–29] followed by recruitment of the histone methyltransferase SETDB1/Egg [30–32]. Upon H3K9 methylation by SETDB1, germline TE insertions may be co-transcriptionally repressed and converted to piRNA generating loci by subsequent recruitment of the HP1 paralog Rhino [25,26,33]. Recruitment of Rhino coincides with non-canonical transcription within the TE insertion and transcripts are directed into a pathway of RNA processing that lacks standard capping, splicing and polyAdenylation [25,34–36]. Since non-canonical transcription can ignore TE encoded termination signals [36] and extend beyond the target TE insertion, transcripts designated for piRNA processing can yield piRNAs from genomic regions outside the TE insertion. This occurs presumably through phased piRNA biogenesis [37–39], since transcripts derived from unique genomic regions flanking the TE will not be the direct target of TE-derived piRNAs that trigger ping-pong biogenesis.

In *Drosophila*, there is evidence that TEs with the capacity to induce flanking H3K9 methylation through piRNA targeting are deleterious due to the silencing of neighboring genes [14,15].

However, there is striking variation in the capacity for TE insertions to trigger these effects. Across two independent strains, only about half of euchromatic insertions show a signature of locally induced H3K9 methylation [15]. Why some TEs trigger local piRNA biogenesis and/or repressive chromatin and others do not is poorly understood, though a variety of factors are known to contribute. One factor is clearly the class of TE. In maize, only some TE families appear to induce local heterochromatin formation [13] and in *Drosophila*, the LTR class appears to exert a stronger effect on local chromatin compared to other families [15]. Such differences may be explained by regulatory sequences embedded within the particular TE family or class.

For example, elements primarily expressed in somatic cells of the ovary trigger a greater degree of flanking H3K9 methylation in cultured ovarian somatic cells [27]. Additionally, TE insertions that lack a promoter and are thus not expressed can fail to trigger flanking piRNA biogenesis in the germline [40].

In the absence of regulatory sequences encoded within TE insertions, the capacity for a TE fragment to nucleate local repression and piRNA biogenesis depends on the interaction between the individual insertion and the transcriptional environment. A recent investigation of flanking piRNA biogenesis triggered by transgenes showed that transcription in opposing directions (convergent transcription) may enhance conversion of TEs into standalone piRNA clusters with flanking piRNA biogenesis [24]. However, there is no general model that explains why some TEs insertions have strong effects on the expression of flanking genes while others do not.

In a genetic analysis of the *Drosophila Mps1* locus, we identified TE insertions that have a complex influence on gene expression whereby insertions can either trigger local gene silencing or de-silencing. Using polyA RNA-seq and small RNA sequencing, we show how the fate of transcripts from this locus shifts between canonical mRNA processing and piRNA biogenesis in the presence of different TE insertions. This complex effect of TE insertions supports a model in which the capacity for one TE to silence flanking genes depends on local patterns of transcription that can be altered by other TE insertions. This represents a case of compensatory mutation or sign epistasis between TE insertions, whereby the harm or benefit of an allele depends on genetic background [41].

## Results

### A DNA transposon insertion rescues a retrotransposon insertion allele of Mps1

The *Drosophila* homolog of *Mps1*, *ald*, is a conserved protein kinase that is a key component of the meiotic and mitotic spindle assembly checkpoint present in most organisms [42–45]. While *Mps1* has both mitotic and meiotic function, the *Drosophila Mps1^A15^* allele only affects meiosis and is caused by a *Doc* non-LTR retrotransposon insertion into the 3’ end of the neighboring gene *alt*, rather than *Mps1* itself [46,47] (Figure 1). *alt* and *Mps1* are convergently transcribed with transcripts overlapping at the 3’ end, a configuration that has been proposed to enable TE insertions to trigger flanking piRNA biogenesis and local gene silencing [24]. To understand why a transposon insertion in one gene could affect the function of another gene, a genetic screen was performed to identify suppressors of the *Doc Mps1^A15^* allele [48]. In this screen, seven stocks were isolated that suppressed nondisjunction caused by the *Mps1^A15^* allele yet retained the *Doc* insertion. Subsequent genetic mapping performed on one of these stocks indicated that the revertant allele was in close proximity to the original *Doc* insertion, so whole genome sequencing of five revertant lines was performed to identify the nature of the revertant lesion. This sequencing revealed no proximal nucleotide differences between the *Mps1^A15^* and revertant alleles, but did identify a new *Hobo* insertion within the flanking gene *alt* (Figure 1). In fact, this insertion was identified in all five revertant stocks, indicating that the identified stocks all carried the same lesion, designated *Mps1^A15.rev^*.

**Figure 1.**
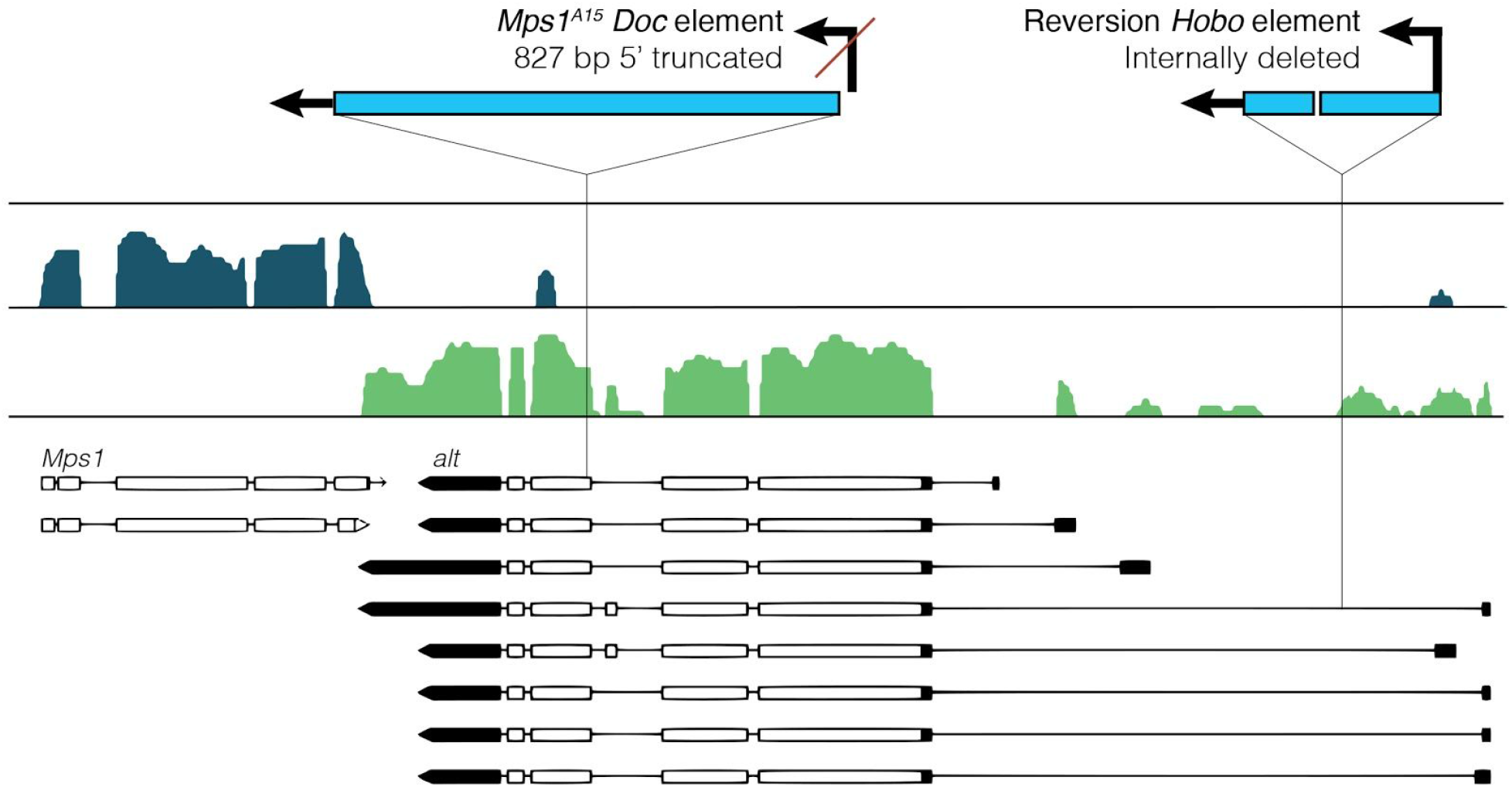
Annotation of TE structure and insertion positions for *Mps1^A15^* and *Mps1^A15.revertant^* alleles within *alt*. Transcript annotations and stranded RNA-seq mappings (blue and green density plots) from ovaries are from Flybase (www.flybase.org) and [50]. The *Mps1^A15^ Doc* insertion is 5’ truncated by 827 bp and located within the 4th/5th exon of *alt*, in the sense direction of the *alt* transcript. The *Mps1^A15.rev^ Hobo* insertion is internally deleted, inserted in the sense direction of *alt* and located in the first intron, 1188 bp form the first *alt* TSS.

PCR and sequencing confirmed the nature of the *Doc* and *Hobo* insertions. The original *Mps1^A15^ Doc* insertion is 5’ truncated and lacks the first 827 nucleotides containing the promoter [49].

The *Doc* is inserted within the fourth of six exons in the sense orientation with respect to *alt*, thus placing a target for germline antisense *Doc* piRNAs within the *alt* transcript. The *Hobo* insertion contains the 5’ and 3’ ends of the consensus *Hobo* element, but is internally deleted. Similar to the *Doc* insertion, it is in the sense orientation with respect to the *alt* transcript, but is inserted within the first intron, 1188 bp from the first TSS. Previous studies indicate that Piwi can repress gene promoter function *via* TE insertions near the TSS [27].

### Local gene silencing by a Doc insertion is ameliorated with insertion of the Hobo element

Since the *Mps1^A15^* allele has an effect on meiosis, but not mitosis, we determined how the two TE insertions influence the germline expression of flanking genes by performing polyA RNA-seq on early embryos. In eggs laid by females homozygous for the *Mps1^A15^* allele, *Mps1* and the neighboring gene *alt* have no expression (Figure 2). Interestingly, the silencing effect of the *Doc* insertion spreads beyond *Mps1* to also cause silencing of *CG7524*, which is divergently transcribed with respect to *Mps1*, while additional genes are not affected. Thus, the *Doc* insertion into *alt* leads to the germline silencing of *alt* and two other genes. RNA-seq on embryos from *Mps1^A15.rev^* homozygous females indicates that the *Hobo* insertion near the 5’ end of *alt* restored the expression of *Mps1* and the flanking gene *CG7524*. However, the silencing of *alt* persists. The *Hobo* insertion does not cause additional silencing of other flanking genes, such as *CG7655*, which is divergently transcribed with respect to *alt* (Figure 2).

**Figure 2.**
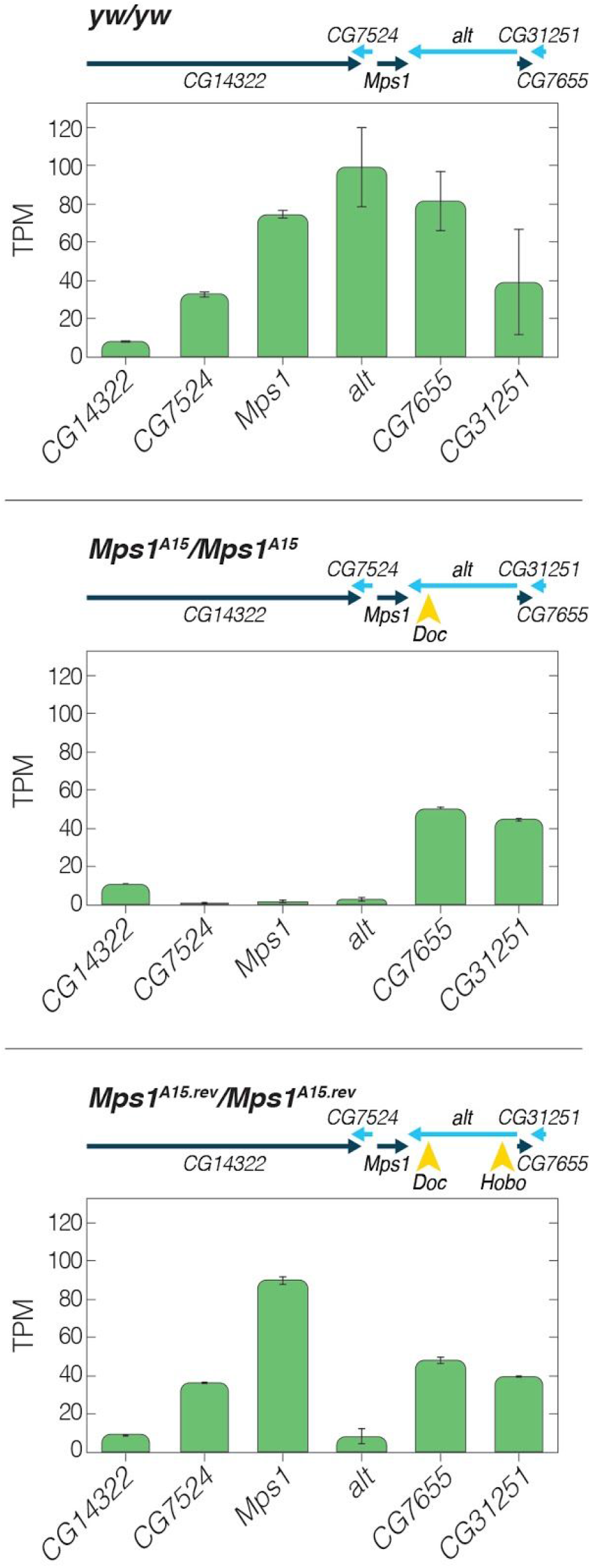
A *Hobo* insertion triggers germline de-silencing of two genes silenced by the *Mps1^A15^ Doc* insertion. PolyA mRNA TPM values from 0-2 hour embryos laid by homozygous mothers. The cartoon above each graph describes which TE insertions (yellow arrowheads) were present in mothers of each experiment. Error bars are S.E. In the presence of only the *Mps1^A15^ Doc* insertion, *CG7524, Mps1* and *alt* are silenced in the germline of *Mps1^A15^* homozygous mothers. In *Mps1^A15.rev^* homozygous mothers, the *Mps1^A15.rev^ Hobo* insertion restores germline expression *CG7524* and *Mps1* in the presence of th *Doc* insertion, but expression of *alt* is not restored.

### Altered gene expression is associated with altered genic piRNA profiles

Since TE insertions have the capacity to induce flanking piRNA biogenesis, we investigated small RNA profiles from whole ovaries in three different experiments. By comparing results with polyA RNA-seq, we would be able to compare modes of transcript processing from this locus, between the standard pathway that includes polyAdenylation with the alternative pathway of piRNA biogenesis that excludes polyAdenylation. Small RNAs were classified as piRNAs based on size (23-30 nt) in both unoxidized and oxidized samples. In the absence of either the *Doc* or *Hobo* insertion, *+/+* ovaries indicate a modest population of piRNAs derived from the site of convergent transcription between *Mps1* and *alt* (Figure 3). Of note, these piRNAs are essentially derived from only one strand, in the sense orientation with respect to *alt* transcription. Across the entire region, there is no evidence that piRNAs are generated through bidirectional transcription since piRNAs derived from one strand do not have a corresponding population derived from the alternate strand. Thus, in the absence of TE insertions, piRNAs from this region appear to be generated through the pathway that generates sense 3’UTR genic piRNAs [51].

**Figure 3.**
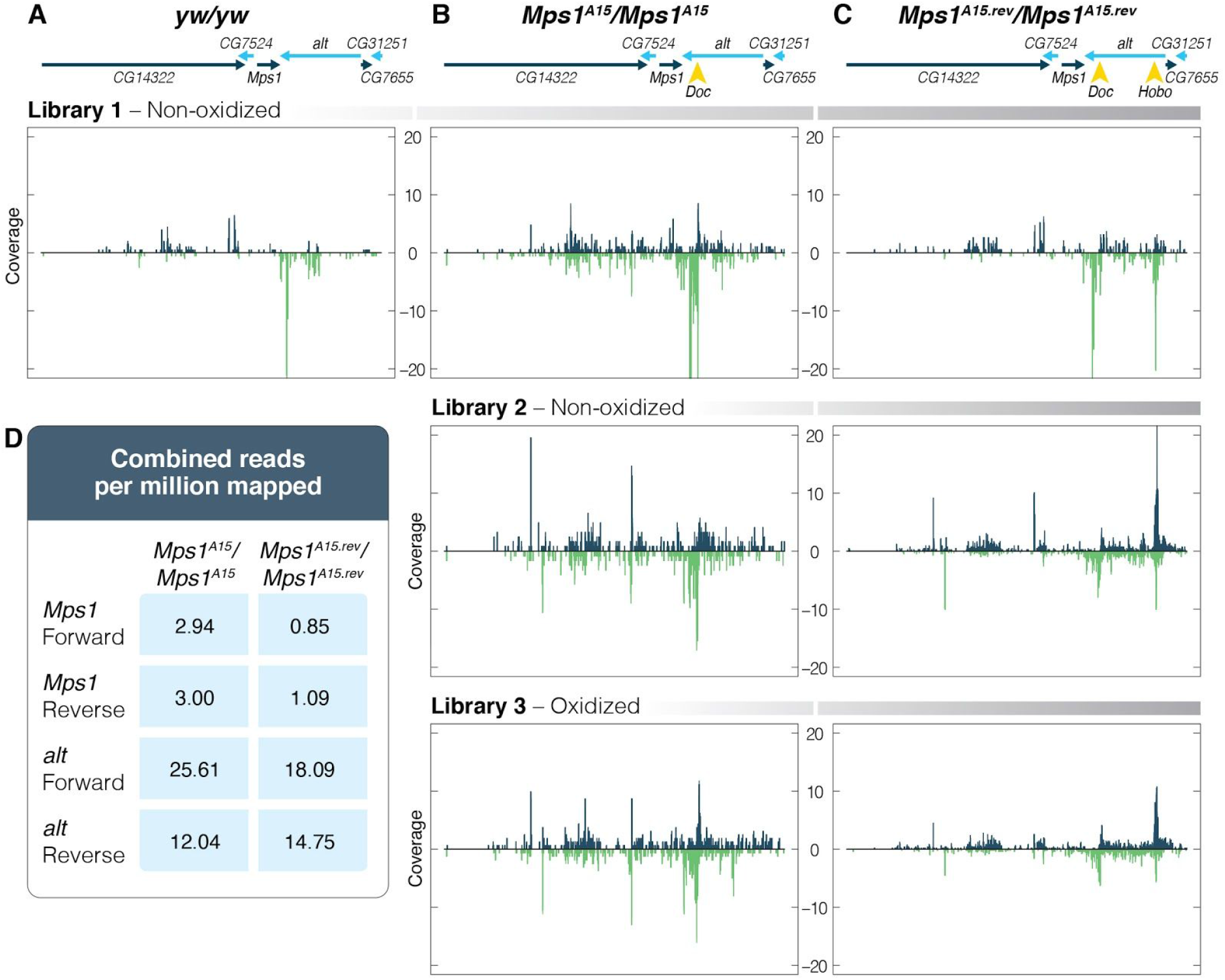
*A Doc* insertion within *alt* triggers the formation of genic piRNA biogenesis from both strands which is disrupted by a *Hobo* insertion. A) Small RNAs derived from convergently transcribed *Mps1* and *alt* are single stranded and in the same direction of the genic transcripts. B) Results across multiple libraries indicate that the *Doc* insertion of the *Mps1^A15^* allele triggers dual-strand piRNA biogenesis across multiple genes. C) The *Hobo* insertion of the *Mps1^A15.rev^* allele disrupts *Doc* triggered dual-strand genic piRNA biogenesis. D) Quantification of mapped reads from combined libraries indicates piRNAs derived from both strands of *Mps1* are reduced by approximately three-fold in *Mps1^A15.rev^* compared to *Mps1^A15^* but maintained at similar levels in both from *alt*.

The insertion of the *Doc* element in *Mps1^A15^* homozygotes changes this pattern dramatically. In all three experiments, the *Doc* insertion is associated with conversion to piRNA biogenesis from both strands. This is evident in *alt*, where the *Doc* insertion is located, but extends across three genes on one side. Interestingly, while piRNAs derived from both strands are identified from *CG14322*, this gene is not silenced, perhaps because the 5’ end of this gene is distant. 23-30 nucleotide RNAs produced from this region have a strong 5’ U bias but do not show a ping-pong signature (Supplemental Figure S1). This supports a model whereby piRNAs are generated through phased-piRNA biogenesis on transcripts generated through bidirectional transcription from within the *Doc* element.

The *Hobo* insertion changes this pattern of flanking piRNA biogenesis. At the site of the *Hobo* insertion near the *alt* TSS, a new population of sense and antisense flanking piRNA emerges. While there is an apparent shift in the location of piRNAs derived from *alt*, the total abundance of *alt* sense and antisense piRNA is similar in *Mps1^A15^* and *Mps1^A15.rev^* homozygotes (Figure 3). However, the *Hobo* insertion is associated with a substantial, though incomplete, reduction of sense and antisense piRNAs derived from flanking genes. In particular, there is an approximate threefold reduction of sense and antisense piRNAs derived from the de-silenced *Mps1*. Overall, while *alt* maintains a population of sense and antisense piRNAs with the *Hobo* insertion, sense and antisense piRNA biogenesis from flanking genes collapses.

### Silencing and de-silencing act zygotically in cis

piRNAs that repress TEs in *Drosophila* are transmitted maternally and maintain continuous silencing across generations [33,52–55]. piRNAs also have the capacity to maintain off-target gene silencing through maternal transmission [56,57]. This maternal transmission also can enable paramutation [58]. Therefore, we tested whether the silencing or de-silencing of *Mps1* depended on the maternal silencing state. This was achieved through quantitative RT-PCR of *Mps1* from ovarian RNA collected from females generated through reciprocal crosses between wildtype, *Mps1^A15^* and *Mps1^A15.rev^* homozygotes. For each of the three pairs of crosses, reciprocal females did not show differences in the expression of *Mps1* (Figure 4). Thus, there is no maternal effect on either silencing or de-silencing. This indicates that piRNAs generated from one allele neither silence nor de-silence in *trans*. Additionally, +/*Mps1^A15.rev^* genotypes have similar expression to wildtype homozygotes and approximately twice the expression level of +/*Mps1^A15^* and *Mps1^A15.rev^/Mps1^A15^* genotypes (Figure 4). Thus, the effect of these alleles on *Mps1* expression appears entirely in *cis*.

**Figure 4.**
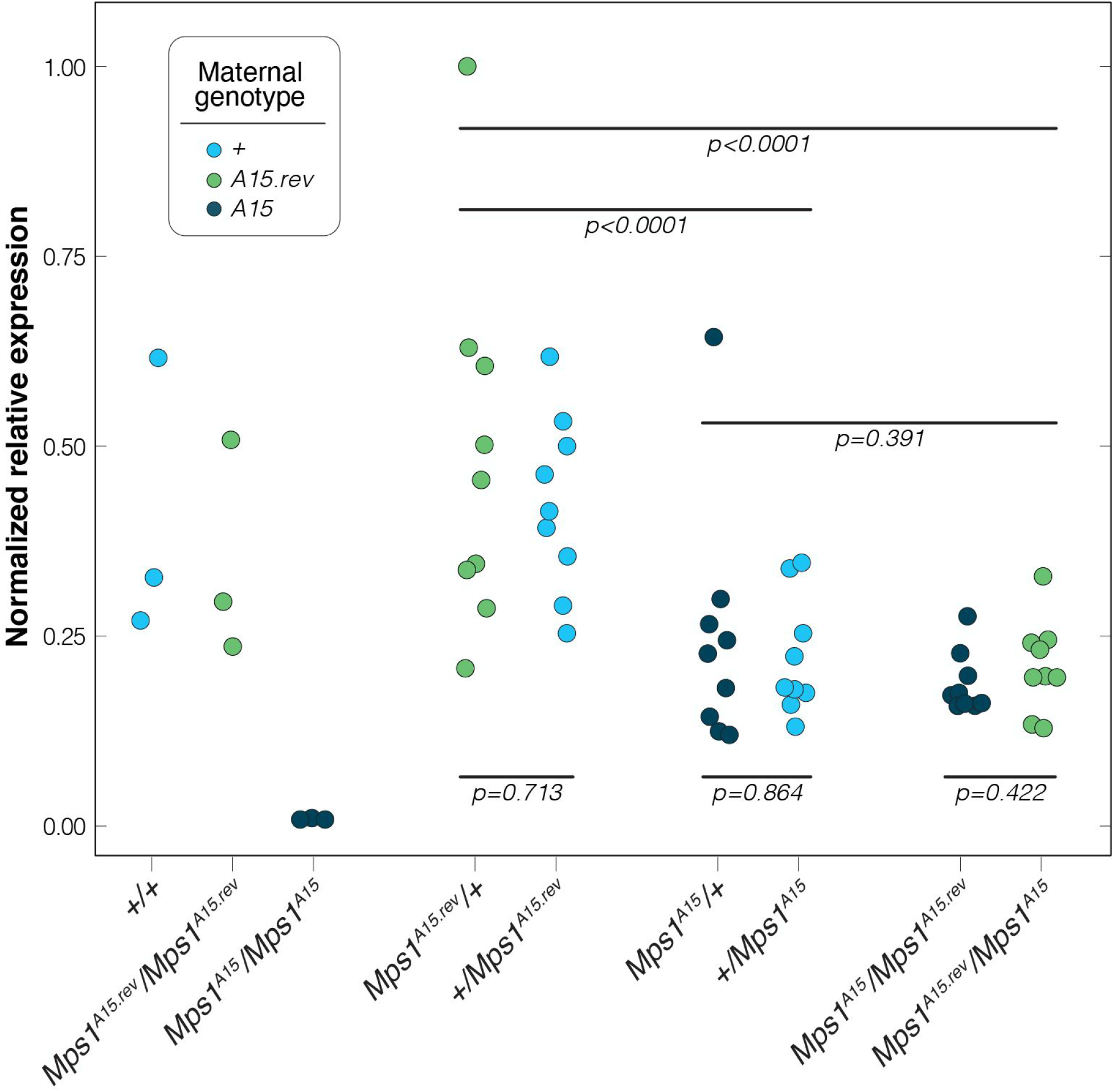
Relative expression (plate normalized) of *Mps1* in ovaries is determined strictly by genotype and the dose of *A15* alleles, indicating a strict zygotic effect on gene expression. Reciprocal progeny of all three pairwise crosses show similar expression levels, indicating no maternal effect. Expression levels of *Mps1* in *Mps1^A15^*/+ and *Mps1^A15^*/*Mps1^A15.rev^* are intermediate between *Mps1^A15^*/*Mps1^A15^* homozygotes and +/+ (or *Mps1^A15.rev^*/*Mps1^A15.rev^*) homozygotes, indicating a strict dose effect of *Mps1^A15^* on *Mps1* expression. Relative expression levels are normalized to maximum 1.0.

## Discussion

Systems of genome defense can maintain genome integrity through the repression of TEs, but off-target effects can lead to gene silencing. These off-target effects have been proposed to contribute to the burden of TEs themselves [5,14]. However, TE insertions near genes do not universally lead to flanking gene silencing. The underlying causes for variation in the effects of TE insertions are not well known. Here we report a pair of TE insertions where one insertion causes gene silencing and the other restores gene expression.

We propose that the original gene silencing of *Mps1* caused by the *Doc* insertion in *alt* can be explained by conversion of the *Doc* element and three neighboring genes (*Mps1, alt* and *CG7524)* into a germline, standalone dual-strand cluster. Even though the truncated *Doc* insertion lacks a promoter, this conversion is likely explained by the insertion of a sense *Doc* target within a transcript driven by the *alt* promoter. The *Doc* fragment in sense orientation likely functions as a target for endogenous antisense *Doc* piRNAs. This conversion is perhaps enhanced by convergent transcription between *Mps1* and *alt* that produces sense piRNAs derived from the 3’ end of *alt* [24]. Upon conversion to a standalone dual-strand cluster, silencing of functional transcripts is likely caused by transcription being directed away from standard mRNA processing into a pathway of piRNA biogenesis [25,34,36]. In this case, flanking piRNA biogenesis from both strands can be considered a readout of genic co-transcriptional repression. How does the *Hobo* insertion lead to de-silencing? The *Hobo* insertion still retains the *Hobo* promoter and TE insertions near gene TSS’s have been shown to block PolII recruitment to genic promoters [27]. Transcriptional repression of *alt* by the *Hobo* insertion thus likely precludes the *Doc* fragment from being a sufficient target for Piwi-piRNAs that can trigger conversion into a piRNA producing locus. Importantly, while the *Hobo* insertion leads to a substantial reduction in the abundance of both sense and antisense *Mps1* piRNAs, flanking piRNA biogenesis is not completely blocked. In this case, one might expect that *Mps1* may retain some repression. Nonetheless, both RNA-seq and RT-qPCR analysis reveal that the *Hobo* insertion completely restores the expression of *Mps1*. This supports the "all-or-nothing" model whereby euchromatic TEs can trigger either weak or strong, but not intermediate, silencing [22,24].

Strikingly, we found no evidence for maternal effects on the expression of *Mps1*, either for silencing alleles or de-silencing alleles. In *Drosophila*, maternal effects by piRNA play an important role in TE repression. This is revealed in syndromes of hybrid dysgenesis where paternal transmission of TEs causes excessive transposition if the mother lacks a corresponding pool of germline piRNAs. Maternal effects in TE regulation also reveal differences in how piRNA source loci depend on piRNAs for either their establishment or maintenance. Functional pericentric dual-strand clusters (such as 42AB) require maternal piRNAs but depletion of Piwi in adult ovaries does not lead to loss of cluster chromatin marks [33] Therefore, dual-strand cluster chromatin can be maintained in the absence of nuclear piRNAs. In contrast, standalone transgenes that trigger flanking piRNA biogenesis require piRNA production for maintenance of *Rhino, HP1* and *H3K9me3* chromatin [24]. Moreover, while maternal inheritance of *I-element* transgenes along with a substantial pool of *I-element* targeting piRNAs can trigger piRNA biogenesis from flanking regions in progeny, this mode of inheritance is not associated with altered chromatin signatures [24]. Overall, it is unclear why maternal effects and paramutation triggered by piRNAs can occur for some genes and not others [56–58].

The costs of gene silencing triggered by TEs have been proposed to shape the dynamics of TEs in populations [5,14]. However, TE insertions do not universally trigger flanking gene repression. In some cases, the expression of neighboring genes can be enhanced [59]. For example, an *Accord* LTR insertion in *Drosophila* melanogaster can enhance the expression of the cytochrome P450 gene *Cyp6g1* and provide resistance to DDT[60]. In *Drosophila simulans*, a *Doc* insertion near *Cyp6g1* has a similar effect and has been the target of positive selection [61]. Enhanced expression is attributed to the regulatory sequences carried by these elements. Here we present a case where TE insertions can alter the germline expression of a gene in opposing ways by altering the local profile of piRNA biogenesis. Formally, this represents a case of sign epistasis. Since the *Hobo* insertion alone is predicted to be deleterious through *alt* silencing, but beneficial when combined with the *Doc* insertion, this satisfies the condition of sign epistasis. For this reason, TE dynamics within populations may not solely be influenced by their single effects, but also their epistatic interactions. As genomic TE density increases, the likelihood for such interactions is expected to increase.

## Materials and Methods

### Identification of the Hobo insertion Mps1^A15^ revertant allele

From an EMS screen [48] to identify suppressors of the *Mps1^A15^ Doc* insertion allele of *ald* seven stocks were identified that suppressed non-disjunction. After one round of recombination with chromosomes carrying P-element *w+* insertions immediately flanking *Mps1* and *alt*, among 351 recombinant chromosomes tested, we were not able to segregate the suppressor lesion away from the *Mps1^A15^* lesion. This indicated that the suppressor lesion was very close to *Mps1* itself. Further analysis revealed that the *Doc* insertion was retained and no nucleotide variants were identified in this region. To identify the nearby suppressor lesion, we performed whole genome sequencing on the original *Mps1^A15^* stock and five of the revertant lines. DNA was prepared from homozygous males or females using the Qiagen DNeasy Blood and Tissue Kit (Samples in Table S1). For each sample 500 ng of DNA was sheared to approximately 600-bp fragments using a Covaris S220 sonicator. KAPA HTP Library Prep Kit for Illumina and Bioo ScientificNEXTflex DNA barcodes were used to prepare libraries which were size selected to 500 – 700 bp. Libraries were quantified using a Bioanalyzer (Agilent Technologies) and a Qubit Fluorometer (Life Technologies). All libraries were pooled and sequenced as 150-bp paired-end samples on an Illumina NextSeq 500 in High-Output mode. Illumina Real Time Analysis version 2.4.11 was run to demultiplex reads and generate FASTQ files.

FASTQ files were aligned to release 6 of the *D. melanogaster* reference genome using bwa version 0.7.7-r441 [62]. SNPs and insertion/deletion polymorphisms were identified using SAMtools and BCFtools (version 0.1.19-44428cd) [63]. Transposable elements were identified as previously described [64]. Briefly, split and discordant read pairs were isolated using SAMBlaster [65] and individual reads were annotated using a BLAST search of the canonical *Drosophila melanogaster* transposable element database [66]. A position with multiple reads from a single TE was defined as a putative TE insertion site and was then manually analyzed. Using this approach, the *Hobo* insertion was identified. Further PCR and Sanger sequencing was used to confirm structure and insertion location within *alt*.

### mRNA Seq

To measure functional gene expression within the germline that is not shunted into piRNA biogenesis, we performed polyA RNA-seq of total RNA collected from 0-2 hour old embryos laid by wildtype, *Mps1^A15^*/*Mps1^A15^* and *Mps1^A15.rev^*/*Mps1^A15.rev^* females. Since zygotic gene expression does not begin until about two hours after egg deposition, this provides a measure of germline gene expression. RNA was obtained from three different collections of pooled embryos per genotype. mRNAseq libraries were generated from 100ng of high-quality total RNA, as assessed using the Bioanalyzer (Agilent). Libraries were made according to the manufacturer’s directions for the TruSeq Stranded mRNA LT Sample Prep Kit – sets A and B (Illumina, Cat. No. RS-122-2101 and RS-122-2102). Resulting short fragment libraries were checked for quality and quantity using the Bioanalyzer (Agilent) and Qubit Fluorometer (Life Technologies).

Libraries were pooled, requantified and sequenced as 75bp paired reads on a high-output flow cell using the Illumina NextSeq instrument. Following sequencing, Illumina Primary Analysis version RTA 2.4.11 and bcl2fastq2 v2.20 were run to demultiplex reads for all libraries and generate FASTQ files. TPM estimates were obtained using the CLC Genomics Workbench.

### Small RNA seq

Small RNA seq was performed using two approaches from RNA collected from whole ovaries of wildtype, *Mps1^A15^*/*Mps1^A15^* and *Mps1^A15.rev^*/*Mps1^A15.rev^* females. One sequencing experiment (Experiment 1) was performed according to the manufacturer’s directions for the TruSeq Small RNA Sample Preparation Kit (Illumina, RS-200-0012). The protocol was adapted to incorporate a 2S blocking DNA oligo for removal of prevalent Drosophila small ribosomal RNA from the sequencing library [67]. Libraries were amplified with 13 PCR cycles and resulting small RNA libraries were cut per the manufacturer’s methods for 20-40 nt cDNA inserts. Short fragment libraries were checked for quality and quantity using the Bioanalyzer and Qubit Fluorometer (Life Technologies). Equal molar libraries were pooled, requantified and sequenced as 75 bp single read on the Illumina NextSeq 500 instrument using NextSeq Control Software 2.2.0.4. At least 6M reads were generated per library, and following sequencing, Illumina Primary Analysis version RTA 2.4.11 and bcl2fastq2 v2.20 were run to demultiplex reads for all libraries and generate FASTQ files.

To ensure piRNAs were being characterized, a second experiment was performed using an altered protocol on oxidized and non-oxidized RNA, in parallel. Pooled RNA samples were split and half the sample was oxidized [68]. RNA from each sample was ligated to 3’ and 5’ adapters using an rRNA blocking procedure [67] and subjected to direct reverse-transcription with unique barcoded RT primers. Barcoded RT products were pooled and size selected on a 10% acrylamide gel for the appropriate size of small RNA cDNAs (18-30 nt) appended to the additional sequence added by the adapters and RT primer. Size selection was facilitated by completing the same procedure in parallel on 18 and 30 nt RNA oligonucleotides. This procedure of pooled size selection allowed all cDNA samples to be extracted under identical conditions. Full protocol is provided in Supplemental. Size selected RT products were extracted from acrylamide, subjected to 15 cycles (non-oxidized) and 18 cycles (oxidized) of PCR and sequenced. Reads were bioinformatically trimmed of adapters, unique molecular identifiers (6bp), size selected between 23 and 30 nt and miRNA/tRNA/rRNA depleted using the CLC Genomic Workbench.

From both experiments, small RNA reads were mapped with bwa to release 6.33 of the *Drosophila melanogaster* reference, counts analyzed with BEDTools [69] and visualized with R. Nucleotide composition and ping-pong signatures were analyzed with the *unitas* package [70].

### Quantitative PCR

RNA was collected from whole ovaries. Sampling was performed in sets of three whereby three daughters were sampled for each of three mothers, thus providing replication across mothers of a given genotype for a total of 9 samples per genotype/mother combination. Total RNA was subjected to Oligo dT reverse transcription (NEB WarmStart RTx) and qPCR performed (NEB Luna qPCR MasterMix) on *ald* and *rp49* (*aldF2: CTG GGC TGC ATC CTT TAC CT; aldR2: TGG CCA TAT GAA CCA GCA TG; rp49F1: ATC GGT TAC GGA TCG AAC AA; rp49R1: GAC AAT CTC CTT GCG CTT CT)*. Each set of three daughters were analyzed on separate plates and statistical analysis was performed using a GLM model (family=gaussian) in R whereby the difference in *ald* and *rp49* Ct values were modeled as a function of plate effects, genotype effects and individual mother effects across cohorts of sisters. No significant effect of the individual mother was found, so this effect was removed from the model. However, a significant effect of plate was identified. Plate values of the difference between *ald* and *rp49* Ct were normalized based on the estimate of this plate effect and further testing was performed using a GLM model for specific comparisons of genotype and to test for maternal effects. Relative expression values are shown normalized to the maximum difference between *ald* and *rp49* Ct values, scaled according to qPCR amplification efficiency of 100%. Primer efficiency values were estimated in real-time and estimated between 98 and 102%

### Data Availability Statement

All sequence data is available at the NCBI under BioProject PRJNA640639. Original data underlying this manuscript can also be accessed from the Stowers Original Data Repository at http://www.stowers.org/research/publications/libpb-XXXX

## Acknowledgements

Many thanks to Jenny Hackett of the KU Genome Sequencing Core for assistance. This core lab is supported by the National Institute of General Medical Sciences (NIGMS) of the National Institutes of Health (NIH) under award number P20GM103638. This project was also supported by NSF MCB Award 1413532 (PI: JPB), NIH-NIGMS/COBRE Award P20GM103638 (PI: Susan Lunte), NIH-NIGMS/AREA Award R15GM099054 (PI: WDG) and NIH-NIGMS/KINBRE Award P20GM103418 (PI: Douglas Wright). Support to MB was provided by the NIH-NIGMS Initiative for Maximizing Student Development program at the University of Kansas (NIH 5R25GM062232). Support to CC was provided by NSF REU Program Award DBI 1560139 (PI: Jennifer Gleason). Additional support was provided by the Stowers Institute for Medical Research and parts of this study were performed in the laboratory of R. Scott Hawley (Stowers Institute for Medical Institute).

---

**Figure S1.**
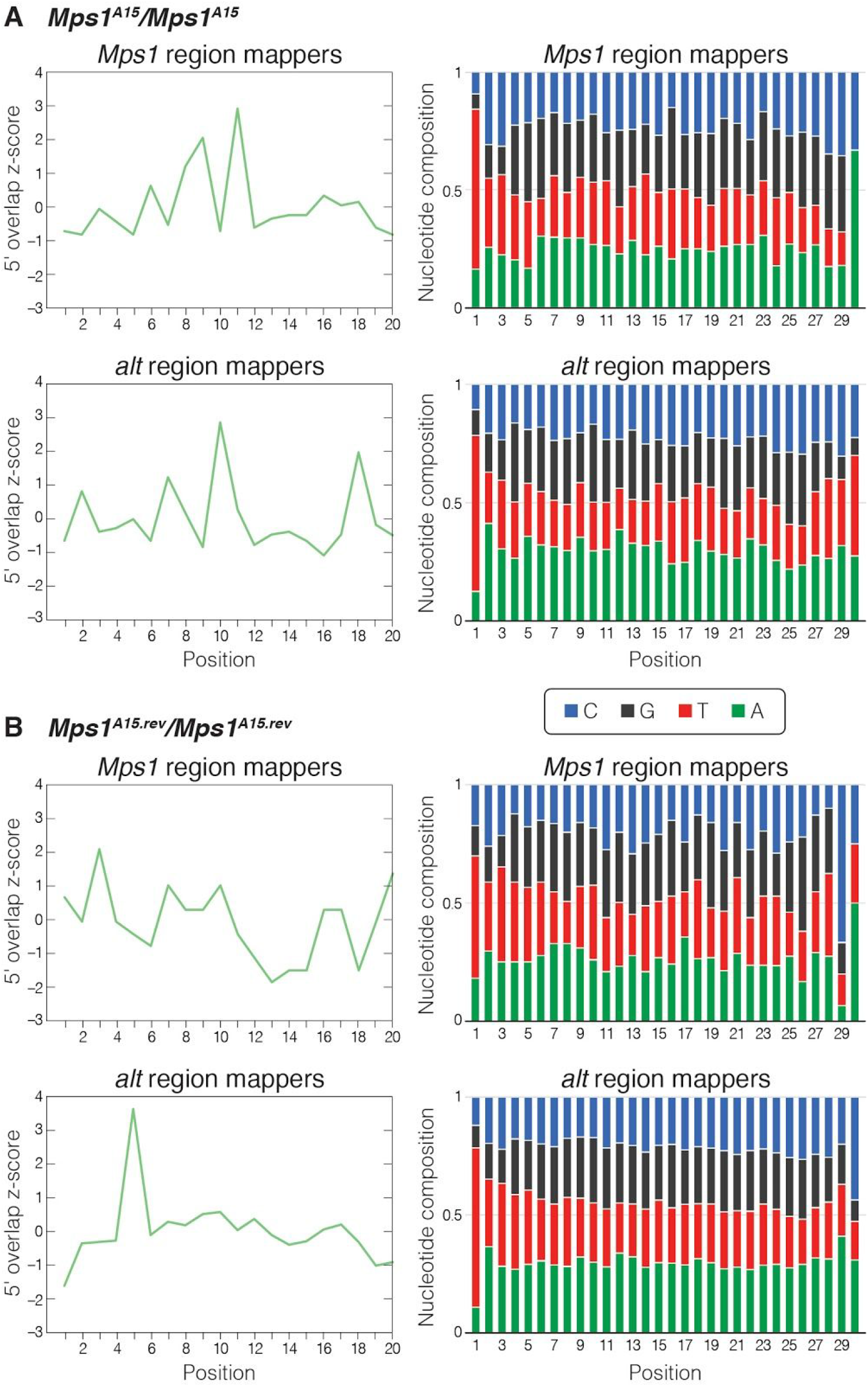
Ping-pong and nucleotide signatures of 23-30 nt small RNAs (from combined libraries) mapping to *Mps1* and *alt* in ovaries of *A15* and *A15*revertant* homozygous females. A U bias is identified as expected for piRNAs. However, no ping-pong signature of 10 bp 5’ overlap is identified among piRNAs mapping to these regions, suggesting instead phased piRNA biogenesis.

